# Deficiency of nucleotide excision repair explains mutational signature observed in cancer

**DOI:** 10.1101/221168

**Authors:** Myrthe Jager, Francis Blokzijl, Ewart Kuijk, Johanna Bertl, Maria Vougioukalaki, Roel Janssen, Nicolle Besselink, Sander Boymans, Joep de Ligt, Jakob Skou Pedersen, Jan Hoeijmakers, Joris Pothof, Ruben van Boxtel, Edwin Cuppen

**Affiliations:** Center for Molecular Medicine and Oncode Institute, University Medical Center Utrecht, Utrecht University, Universiteitsweg 100, 3584, CG, Utrecht, The Netherlands; Department of Molecular Medicine, Aarhus University, Palle Juul-Jensens Boulevard 99, 8200 Aarhus N, Denmark; Erasmus Medical Center, Wytemaweg 80, 3015 CN Rotterdam, The Netherlands

**Author notes:** These authors contributed equally to this work (Myrthe Jager, Francis Blokzijl). Present address: Princess Máxima Center for Pediatric Oncology, 3584 CT Utrecht, The Netherlands.

**Keywords:** ERCC1, XPC, nucleotide excision repair, NER, adult stem cell, liver, small intestine, organoids, mutational signatures, Signature 8, cancer, progeria, CRISPR-Cas

## Abstract

Nucleotide excision repair (NER) is one of the main DNA repair pathways that protect cells against genomic damage. Disruption of this pathway can contribute to the development of cancer and accelerate aging. Tumors deficient in NER are more sensitive to cisplatin treatment. Characterization of the mutational consequences of NER-deficiency may therefore provide important diagnostic opportunities. Here, we analyzed the somatic mutational profiles of adult stem cells (ASCs) from NER-deficient *Ercc1^-/Δ^* mice, using whole-genome sequencing analysis of clonally derived organoid cultures. Our results indicate that NER-deficiency increases the base substitution load in liver, but not in small intestinal ASCs, which coincides with a tissue-specific aging-pathology observed in these mice. The mutational landscape changes as a result of NER-deficiency in ASCs of both tissues and shows an increased contribution of Signature 8 mutations, which is a pattern with unknown etiology that is recurrently observed in various cancer types. The scattered genomic distribution of the acquired base substitutions indicates that deficiency of global-genome NER (GG-NER) is responsible for the altered mutational landscape. In line with this, we observed increased Signature 8 mutations in a GG-NER-deficient human organoid culture in which *XPC* was deleted using CRISPR-Cas9 gene-editing. Furthermore, genomes of NER-deficient breast tumors show an increased contribution of Signature 8 mutations compared with NER-proficient tumors. Elevated levels of Signature 8 mutations may therefore serve as a biomarker for NER-deficiency and could improve personalized cancer treatment strategies.

## INTRODUCTION

The genome is continuously exposed to mutagenic processes, which can damage the DNA and can ultimately result in the accumulation of mutations. To counteract these processes, cells exploit multiple DNA repair pathways that each repair specific lesions. Deficiency of these pathways can contribute to cancer initiation and progression. To increase insight into the cellular processes that underlie mutation accumulation, such as deficiency of specific DNA repair pathways, genome-wide mutational patterns of tumors can be characterized (Alexandrov et al. 2013; Nik-Zainal et al. 2016). To date, systematic analyses of tumor genomes have revealed 30 signatures of base substitutions and 6 rearrangement signatures of mutational processes in cancer genomes (Alexandrov et al. 2013; Nik-Zainal et al. 2016). These mutational signatures may have important diagnostic value. For example, several signatures are associated with BRCA1/2 inactivity and can consequently be predictive for a response to PARP inhibition or cisplatin treatment (Waddell et al. 2015; Davies et al. 2017).

Although for some signatures the underlying molecular process (Kim et al. 2016; Alexandrov et al. 2013, 2016) or involved DNA repair pathway (Kim et al. 2016; Davies et al. 2017; Alexandrov et al. 2013) is known, in-depth mechanistic insight is still lacking for the majority of the mutational signatures (Petljak and Alexandrov 2016). Efforts to link mutational processes to specific signatures have mainly focused on associating mutation data from tumors to mutagen exposure and DNA repair-deficiency. Yet, tumors are genomically highly unstable and typically multiple processes have contributed to mutation accumulation (Alexandrov et al. 2013; Nik-Zainal et al. 2016), which hampers the identification of the processes that cause specific mutational signatures. We recently developed an approach for measuring mutations in non-cancerous adult stem cells (ASCs), by combining organoid culturing technology with whole-genome sequencing (WGS) (Jager et al. 2018; Drost et al. 2017). This method can be used to determine the mutations that have accumulated during life and during culturing. Tissue-specific ASCs maintain a highly stable genome both *in vivo* and *in vitro,* and therefore provide a stable system to study mutational processes in detail (Blokzijl et al. 2016; Huch et al. 2015). Furthermore, ASCs constitute a relevant cell source to study mutational patterns, as these cells are believed to be the cell-of-origin for specific types of cancer (Barker et al. 2009; Zhu et al. 2016; Adams et al. 2015).

Using this technique, we set out to determine the mutational consequences of deficiency of nucleotide excision repair (NER). NER is one of the main cellular DNA repair pathways (lyama and Wilson 2013), and consists of two subpathways: global-genome NER (GG-NER), which repairs bulky helix-distorting lesions throughout the genome, and transcription-coupled NER (TC-NER), which resolves RNA polymerase blocking lesions during transcription (lyama and Wilson 2013; Marteijn et al. 2014; Hoeijmakers 2009). Somatic mutations in *ERCC2,* a key factor of NER, were previously associated with Signature 5 in urothelial tumors (Kim et al. 2016). However, NER has been suggested to underlie multiple mutational signatures, based on large-scale tumor mutation analyses (Alexandrov et al. 2013), and not all NER-deficient tumors are characterized by a high Signature 5 contribution (Kim et al. 2016). This observation suggests that NER-deficiency might be associated with other mutational signatures as well.

To characterize the mutational consequences of NER-deficiency, we studied mutagenesis in *Ercc1^-/Δ^* mice and XPC-knockout (*XPC*^KO^) organoids. ERCC1 plays a crucial role in the core NER pathway involving both GG-NER and TC-NER (Kirschner and Melton 2010; Iyama and Wilson 2013; Sijbers et al. 1996a; Aboussekhra et al. 1995), in crosslink repair (Rahn et al. 2010), and in single strand annealing (SSA) of double strand breaks (Al-Minawi et al. 2008). *ERCC1* is mutated in ~4.5% of all human tumors, especially skin and liver cancer (http://dcc.icgc.org), and single nucleotide polymorphisms in *ERCC1* have been linked to an increased risk of developing colorectal cancer (Ni et al. 2014). *Ercc1^-/Δ^* mice are hemizygous for a single truncated *Ercc1* allele, which largely corrupts protein function (Dollé et al. 2011; Weeda et al. 1997) and results in decreased NER-activity (Su et al. 2012). *Ercc1^-/Δ^* mice have a reduced lifespan as a result of progeroid-like symptoms and live five times shorter than wild-type (WT) littermates (Dollé et al. 2011; Vermeij et al. 2016). The livers of *Ercc1^-/Δ^* mice display various aging-like characteristics and pathology (Dollé et al. 2011; Gregg et al. 2012; Niedernhofer et al. 2006; Weeda et al. 1997), whereas, other organs, such as the small intestine, do not show an obvious pathological phenotype. Thus the consequences of loss of ERCC1 differ considerably between tissues, although the reason for this remains unclear. XPC is involved in the recognition of bulky DNA adducts in the GG-NER pathway specifically (Puumalainen et al. 2015; Iyama and Wilson 2013). Germline mutations in this gene cause Xeroderma Pigmentosum, a disorder characterized by enhanced sensitivity to UV-light and development of various cancer types at an early age (Sands et al. 1995; Melis et al. 2008; Dupuy and Sarasin 2015).

Here, we combined the organoid culture system with *in vivo* and *in vitro* knockout models, providing the unique opportunity to characterize the genome-wide mutational consequences of NER-deficiency in a stable genetic background. Furthermore, we compared the genome-wide mutational patterns of NER-deficient and NER-proficient tumors from a breast cancer cohort (Nik-Zainal et al. 2016). Both quantitative and qualitative mutational differences that are associated with NER status were identified, creating novel insight into the molecular processes underlying mutation accumulation, cancer, and aging.

## RESULTS

### Loss of NER protein ERCC1 increases the number of base substitutions in liver, but not in small intestinal mouse ASCs

To characterize the mutational consequences of NER-deficiency, we generated clonal organoid cultures from single liver and small intestinal ASCs of three female *Ercc1^-/Δ^* mice and three female WT littermates (Fig. 1A). The tissues were harvested at the age of 15 weeks, which is the time point at which *Ercc1^-/Δ^* mice generally start to die as a consequence of early aging pathologies (Vermeij et al. 2016). WGS analysis of DNA isolated from the clonal organoid cultures allows for reliable determination of the somatic mutations that were accumulated during life in the original ASCs (Blokzijl et al. 2016; Jager et al. 2018). Subclonal mutations acquired after the single-cell-step will only be present in a subpopulation of the cells and are filtered out based on a low allele frequency (Jager et al. 2018). We also sequenced the genomes of polyclonal control biopsies from the tail of each mouse, which served as control samples to exclude germline variants.

**Figure 1.**
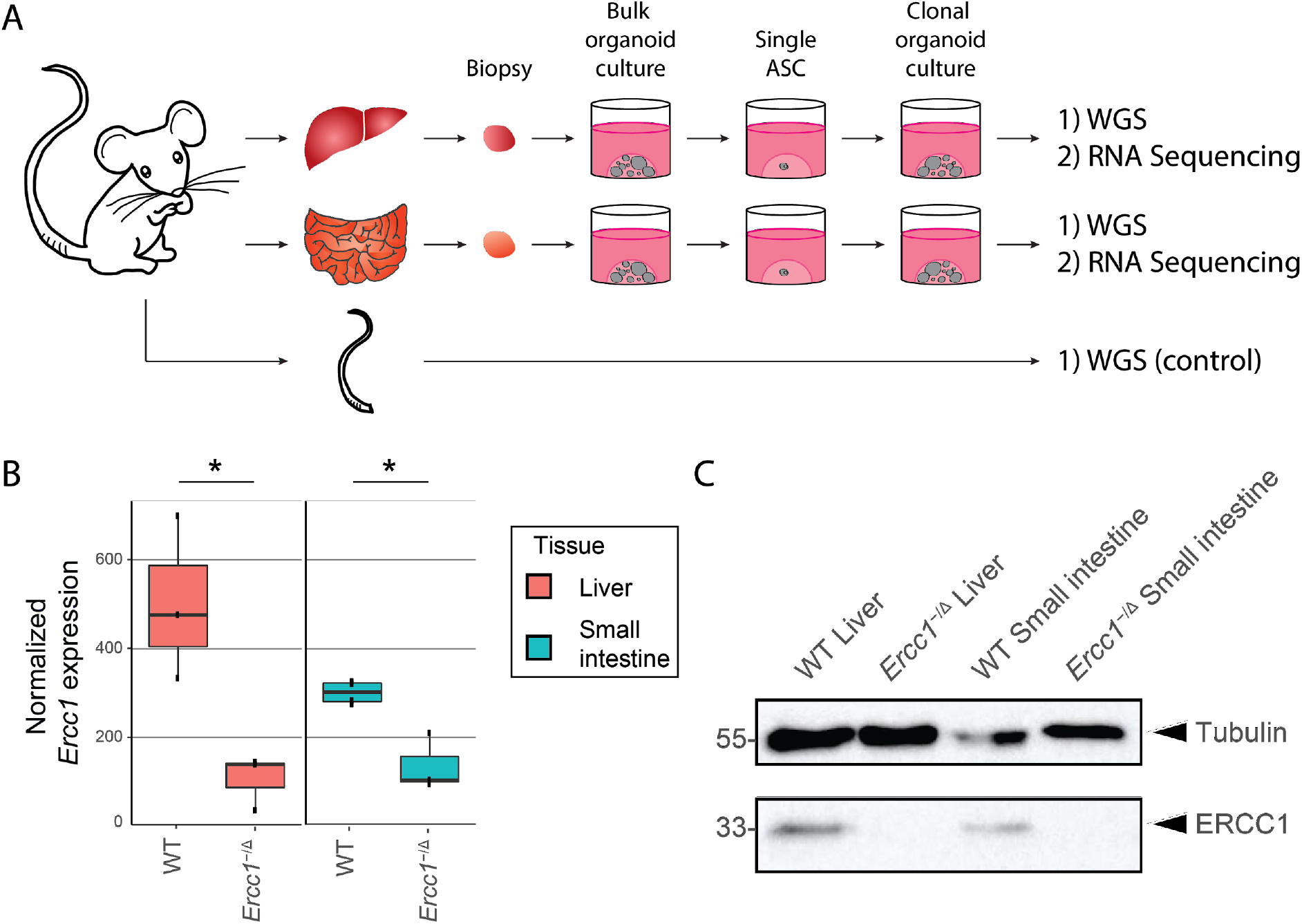
Experimental setup and tissue-specific expression of *Ercc1* in mouse ASCs. (A) Schematic overview of the experimental setup used to determine the mutational patterns in single ASCs from the liver and small intestine of mice. Biopsies from the liver and small intestine of six 15-week-old female mice (three *Ercc1^-/Δ^* mice and three WT littermates) were cultured in bulk for ~1.5 week to enrich for ASCs. Subsequently, clonal organoids were derived from these bulk organoid cultures and expanded for approximately 1 month, until there were enough cells to perform both WGS and RNA sequencing. As a control sample for filtering germline variants, a biopsy of the tail of each mouse was also subjected to WGS. (B) Boxplots depicting normalized *Ercc1* expression in ASC organoid cultures from liver and small intestine of *Ercc1^-/Δ^* mice (n = 3 and n = 3, respectively) and WT littermates (n = 3 and n = 4, respectively). Asterisks represent significant differences (P < 0.05, negative binomial test). (C) Western blot analysis of ERCC1 in *Ercc1^-/Δ^* and WT small intestinal and liver mouse organoids.

To determine transcriptome profiles, we performed RNA sequencing on one clonal organoid culture from each tissue of each mouse. *Ercc1* is significantly differentially expressed (P < 0.05, negative binomial test) between WT and *Ercc1^-/Δ^* in both liver and small intestinal ASCs (Fig. 1B), confirming the anticipated effects of the *Ercc1* mutations at the mRNA level. While there is some *Ercc1* expression in *Ercc1^-/Δ^* ASCs, the C-terminal domain of ERCC1 is essential in ERCC1-XPF complex formation and disruption of this interaction reduces the stability of ERCC1 protein (Tripsianes et al. 2005; de Laat 1998; Sijbers et al. 1996b). Indeed, ERCC1 protein is not detectable by immunoblotting in *Ercc1^-/Δ^* organoid cultures of both tissues (Fig. 1C). No other DNA repair genes were differentially expressed between WT and *Ercc1^-/Δ^* ASCs (Supplemental File S1). Notably, the expression of 8 out of 9 core NER genes, including *Ercc1,* is higher in WT liver ASCs than WT small intestinal ASCs (Supplemental Fig. S1, Supplemental Table S1).

WGS analysis on the clonally-expanded organoid cultures revealed 4,238 somatic base substitutions in the autosomal genome of 11 clonal ASC samples (Fig. 2A; Supplemental Table S2). Targeted deep-sequencing validated ~97.5% of these base substitutions (Supplemental File S2). Liver ASCs of WT mice acquired 19.5 ± 4.1 (mean ± standard deviation) base substitutions per week. This rate is similar in ASCs of the small intestine, at 16.1 ± 3.1 mutations per week, and is in line with the observation that human liver and intestinal ASCs have similar mutation accumulation rates *in vivo* (Blokzijl et al. 2016). Loss of ERCC1 induced a twofold increase (45.5 ± 3.0 base substitutions per week) in the number of base substitutions in ASCs of the liver (Fig. 2A, Supplemental Fig. S2A). However, we did not observe a different mutation rate in small intestinal ASCs of *Ercc1^-/Δ^* mice (21.0 ± 4.9 base substitutions per week) compared with WT small intestinal ASCs (Fig. 2A, Supplemental Fig. S2A). We also observed a significant increase in the number of double base substitutions in liver ASCs lacking ERCC1 (*q* < 0.05, ř-test, FDR correction; Fig. 2B, Supplemental Fig. S2B, Supplemental Table S3). *Ercc1^-/Δ^* liver ASCs acquire 0.49 ± 0.06 double base substitutions per week, while WT liver ASCs acquire only 0.05 ± 0.04 double base substitutions per week. Again, we did not observe this difference between WT and mutant ASCs of the small intestine (0.07 ± 0.10 and 0.07 ± 0.07 per week, respectively). The increased number of double base substitutions in the liver ASCs remained significant after normalizing for the total number of base substitutions (q < 0.05, ř-test, FDR correction; Supplemental Fig. S2C), indicating a liver-specific enrichment of double base substitutions in *Ercc1^-/Δ^* ASCs compared with WT.

**Figure 2.**
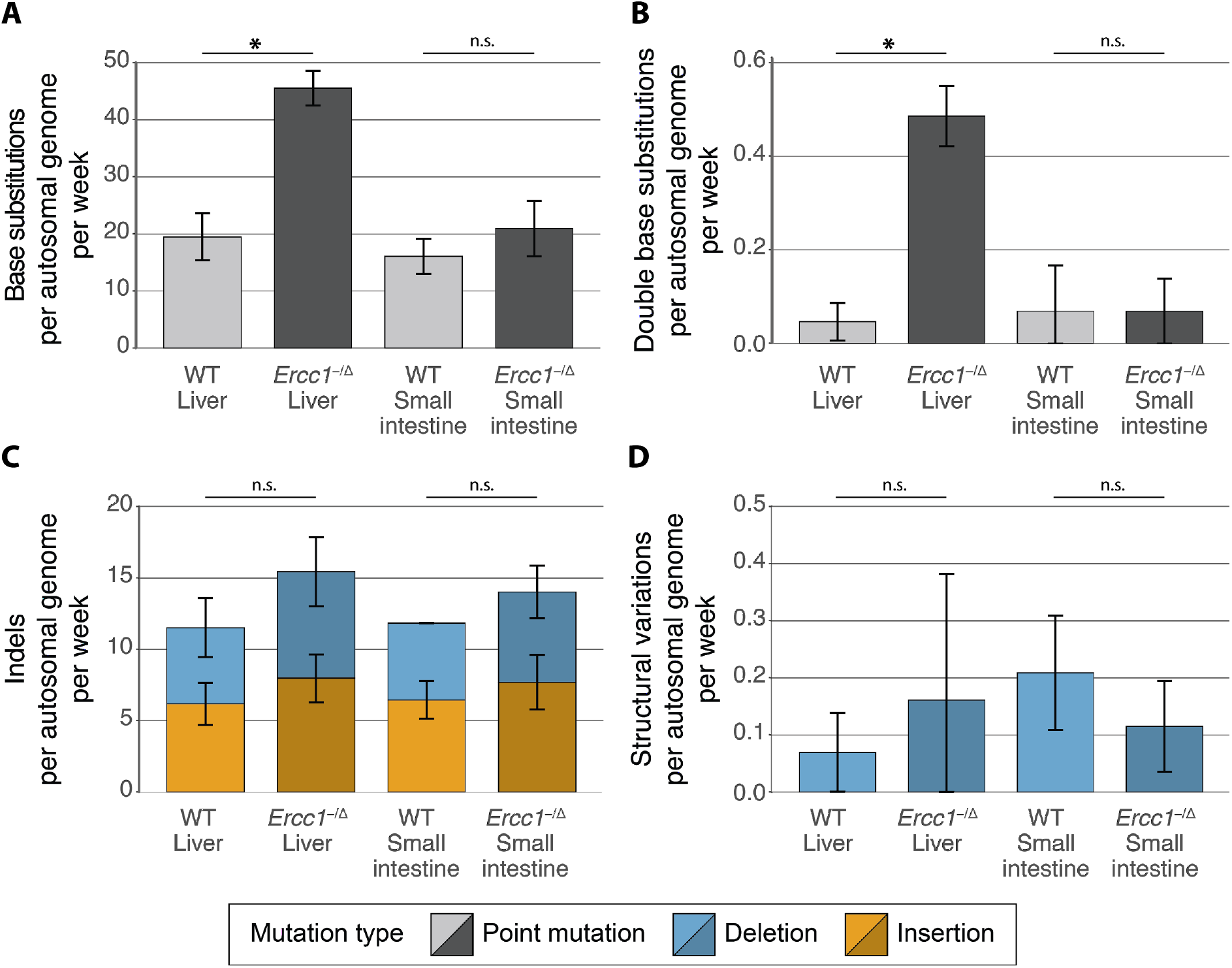
Somatic mutation rates in the genomes of ASCs from liver and small intestine of WT and *Ercc1^-/Δ^* mice. (A) Base substitutions, (B) double base substitutions, (C) indels, and (D) SVs acquired per autosomal genome per week in ASCs of WT liver (n = 3), *Ercc1^-/Δ^* liver (n = 3), WT small intestine (n = 2), and *Ercc1^-/Δ^* small intestine (n = 3). Error bars represent standard deviations. Asterisks represent significant differences (*q* < 0.05, two-sided t-test, FDR correction). n.s. : non-significant (q ≥ 0.05, two-sided t-test, FDR correction).

In addition to the 4,238 base substitutions, we identified 2,116 small insertions and deletions (indels) and 21 larger deletions (≥100 bp) in the autosomal genome of the 11 clonal ASC samples (Supplemental Table S2). As opposed to the base substitutions, we observed similar indel numbers for WT and *Ercc1^-/Δ^* ASCs of both tissues (Fig. 2C, Supplemental Fig. S2D). Of note, accurate identification of indels is more challenging than base substitutions, and as a result, these calls may contain more false positives. ASCs in the small intestine and liver of the mice acquire approximately 13.3 ± 3.4 indels per week, independent of *Ercc1* mutation status. Likewise, loss of ERCC1 did not influence the number or type of structural variations (SVs) in ASCs of the small intestine and the liver (Fig. 2D, Supplemental Fig. S2E, Supplemental Table S2). Each mouse ASCs carried 0 - 6 deletions (median length of 539 bp; Supplemental Table S4). Finally, a genome-wide copy-number profile was generated to identify chromosomal gains and losses. These profiles indicated that all WT and *Ercc1^-/Δ^* ASCs were karyotypically stable during life (Supplemental Fig. S3). Nevertheless, some subclonal aneuploidies were detected in a WT as well as an *Ercc1^-/Δ^* liver organoid sample, which are most likely culturing artefacts that occurred *in viřro* after the clonal step irrespective of *Ercc1* mutation status.

### Loss of NER protein ERCC1 induces Signature 8 mutations in mouse ASCs

To further dissect the mutational consequences of NER-deficiency, we characterized the mutation spectra in the mouse ASCs. Regardless of tissue-type, the mutation spectra of all assessed ASCs are predominantly characterized by C:G > A:T mutations and C:G > T:A mutations (Fig. 3A). However, the mutation spectra of NER-proficient and NER-deficient ASCs differed significantly for both tissues (q < 0.05, X^2^-test, FDR correction). Indeed, there are some notable differences, such as an increased contribution of T:A > A:T mutations in *Ercc1^-/Δ^* ASCs compared with WT ASCs (Fig. 3A).

**Figure 3.**
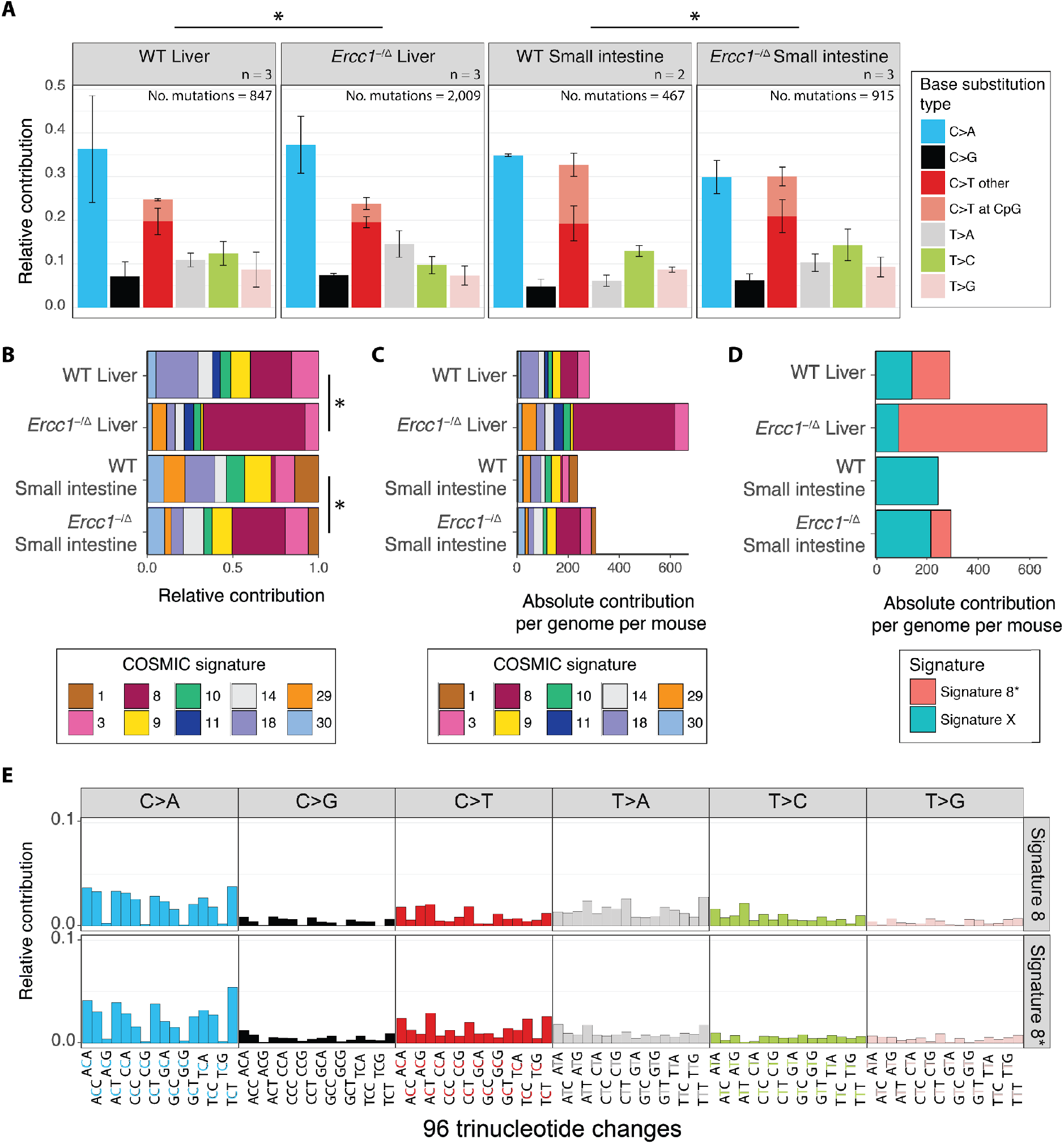
Mutational patterns of base substitutions acquired in the genomes of ASCs from liver and small intestine of WT and *Ercc1^-/Δ^* mice. (A) Mean relative contribution of the indicated mutation types to the mutation spectrum for each mouse ASC group. Error bars represent standard deviations. The total number of mutations, and total number of ASCs (n) per group is indicated. Asterisks indicate significant differences in mutation spectra (q < 0.05, X^2^-test, FDR correction). (B) Relative contribution of the indicated COSMIC mutational signatures to the average 96-channel mutational profiles of each mouse ASC group. Asterisks indicate significantly different signature contributions, *P* values where obtained using a bootstrap resampling approach (Methods, Supplemental Fig. S6E-F) (C) Absolute contribution of the indicated COSMIC mutational signatures to the average 96-channel mutational profiles of each mouse ASC group. (D) Absolute contribution of two mutational signatures that were identified by non-negative matrix factorization (NMF) analysis of the average 96-channel mutational profiles of each mouse ASC group. (E) Relative contribution of each indicated context-dependent base substitution type to mutational Signature 8 and Signature 8*.

To gain insight into these differences, we generated 96-channel mutational profiles of all ASCs (Supplemental Fig. S4, Supplemental Fig. S5) and assessed the contribution of each COSMIC mutational signature (http://cancer.sanger.ac.uk/cosmic/signatures) to the average 96-channel mutational profile (centroid) per group (Supplemental Fig. S6B). We could reconstruct the original centroids well with the 30 COSMIC signatures, as the reconstructed centroids are highly similar to the original centroids for all four ASC groups (average cosine similarity = 0.95, Supplemental Fig. S6A). The contribution of the COSMIC signatures is significantly different between NER-proficient and NER-deficient ASC groups for both liver and small intestine (*d > d_WT_0.05_ and d > d_MuT__0.05,* bootstrap resampling method, see Methods, Supplemental Fig. S6C-D).

We subsequently reconstructed the 96-channel mutational profiles using the top 10 contributing COSMIC mutational signatures (Fig. 3B-C). We could reconstruct the centroids comparably well using this subset of 10 COSMIC signatures (average cosine similarity = 0.95, Supplemental Fig. S6A). The 96-channel mutational profiles of NER-deficient liver ASCs not only closely resemble Signature 8 (cosine similarity of 0.92; Supplemental Fig. S7), but Signature 8 can almost fully explain the increase in base substitutions in NER-deficient liver ASCs (Fig. 3C). The number of Signature 8 mutations is also increased in all small intestinal ASCs of *Ercc1^-/Δ^* mice compared with WT small intestinal ASCs (Fig. 3C). This finding shows that NER-deficiency can result in elevated numbers of Signature 8 mutations in ASCs, regardless of tissue-type.

In addition, we performed an unbiased signature analysis by extracting two mutational signatures *de novo* from the mouse mutation catalogs using non-negative matrix factorization (NMF) (Supplemental File S3, Supplemental Fig. S8). One of the identified signatures, Signature X, contributes approximately 100 mutations to liver ASCs and 200 mutations to small intestinal ASCs, in both WT and *Ercc1^-/Δ^* mice (Fig. 3D), suggesting that this signature represents a mutational process that is generally active in mouse ASCs. In line with this, Signature X is highly similar to 96-channel mutational profiles of ASCs of the small intestine of old mice (Behjati et al. 2014) (cosine similarity = 0.95, Supplemental Fig. S8B). As expected, this mouse signature is not similar to any of the known COSMIC signatures identified in human tumor sequencing data (Supplemental Fig. S8B). The other signature, Signature 8*, is highly similar to COSMIC Signature 8 (cosine similarity = 0.91; Fig. 3E, Supplemental Fig. S8B) and has an increased contribution in *Ercc1^-/Δ^* liver ASCs compared with WT (Fig. 3D; Supplemental Fig. S8C). Moreover, the contribution of Signature 8* mutations is also increased in *Ercc1^-/Δ^* small intestinal ASCs in comparison to WT small intestinal ASCs (Fig. 3D; Supplemental Fig. S8C). These findings confirmed that NER-deficiency results in base substitutions that show a 96-channel profile similar to COSMIC Signature 8.

Mutations are distributed non-randomly throughout the genome in cancer cells and in human ASCs (Schuster-Böckler and Lehner 2012; Blokzijl et al. 2016). NER is one of the pathways that is suggested to underlie this non-random distribution of mutations (Perera et al. 2016; Zheng et al. 2014). Firstly, NER-activity has been linked to a local enrichment of mutations at gene promoters (Perera et al. 2016). However, we do not observe any significant differences in the depletion of mutations in promoters, promoter-flanking, and enhancer regions between NER-proficient and -deficient ASCs (Supplemental Fig. S9A). Secondly, TC-NER results in a depletion of mutations in expressed genes, as this pathway repairs lesions on the transcribed strand during transcription (Pleasance et al. 2010). Mutations are indeed depleted in genic regions of NER-proficient WT mouse ASCs, but the depletion is not significantly different in NER-deficient ASCs (n.s., Poisson test, FDR correction; Supplemental Fig. S9A). Moreover, the average expression levels of genes in which the somatic mutations are located do not differ between *Ercc1^-/Δ^* and WT ASCs (n.s., ř-test, FDR correction; Supplemental Fig. S9B), suggesting that *Ercc1^-/Δ^* ASCs do not accumulate more mutations in expressed genes. Finally, there are no obvious changes in transcriptional strand bias, although the mutation numbers are too low to be conclusive (Supplemental Fig. S9C). NER-deficiency thus influences both the mutation load and mutation type, but not the genomic distribution of the observed base substitutions in mouse ASCs, suggesting that the contribution of TC-NER in the observed mutational consequences is minimal in these cells.

### Loss of GG-NER protein XPC induces Signature 8 mutations in human ASCs

To identify a potential causal relationship between NER-deficiency and Signature 8 in human ASCs, we generated a human GG-NER deficient *XPC*^KO^ ASC using CRISPR-Cas9 gene-editing in a human small intestinal organoid culture (Fig. 4A). After confirming absence of XPC protein (Fig. 4B), we passaged the *XPC*^KO^ clone for approximately 2 months to allow the accumulation of sufficient mutations for downstream analyses. Subsequently, we derived subclonal cultures of single ASCs and expanded these until sufficient DNA could be isolated for WGS. This approach allowed us to catalog the mutations that specifically accumulated between the two clonal expansion steps in the absence of XPC (Supplemental Fig. S10A) (Drost et al. 2017; Blokzijl et al. 2016; Jager et al. 2018). As a control, WGS data of three previously-established XPC^WT^ organoid cultures of the same human donor was used (Blokzijl et al. 2016).

**Figure 4.**
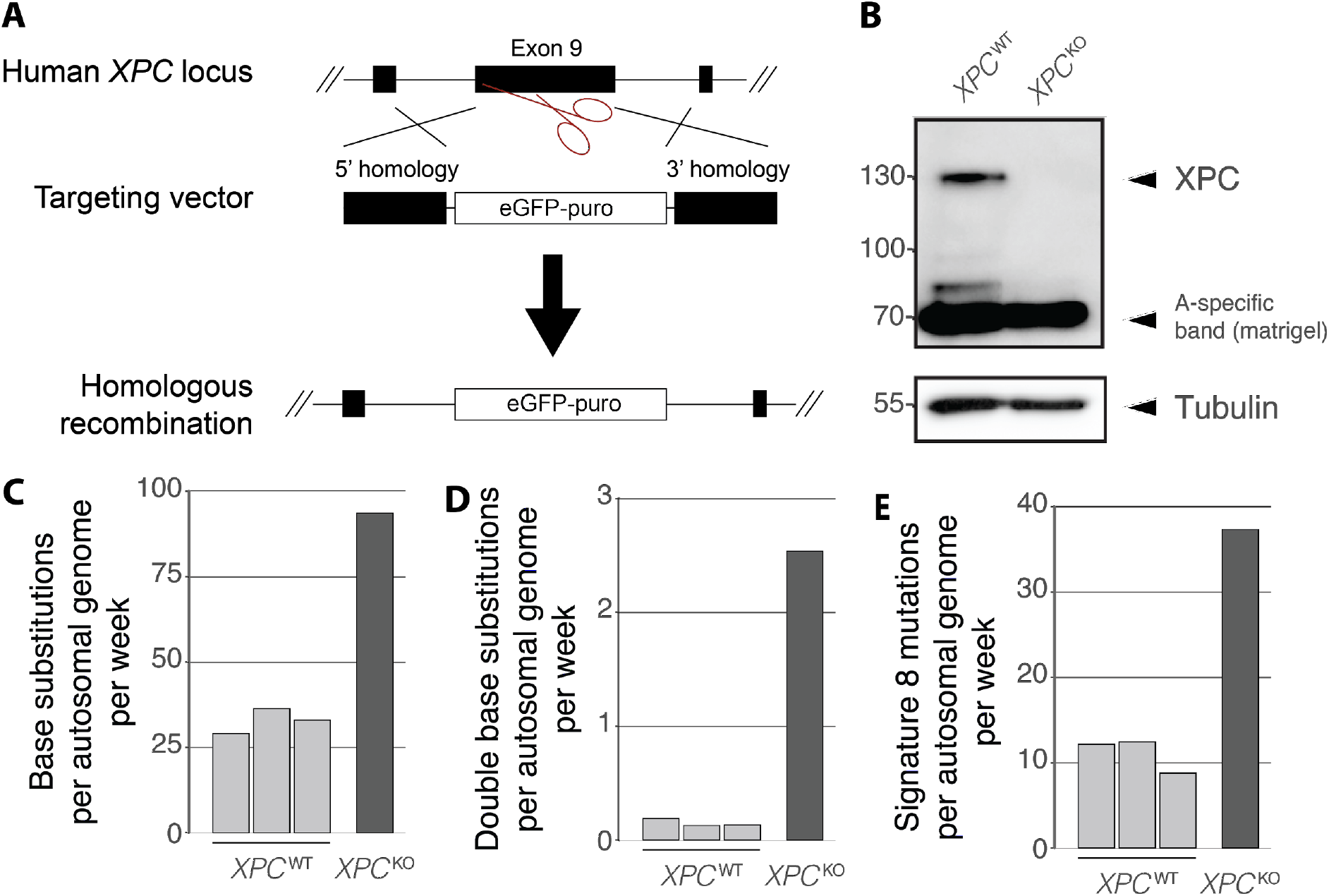
Mutational consequences of *XPC*^KO^ in human intestinal organoid cultures *in vitro.* (A) Targeting strategy for the generation of *XPC*^KO^ organoid cultures using CRISPR-Cas9 gene-editing. (B) Western blot analysis of XPC in human *XPCA*^WT^ and *XPC*^KO^ organoids. (C) Number of base substitutions, (D) double base substitutions, and (E) Signature 8 mutations acquired per autosomal genome per week in human *XPCA*^WT^ ASCs (n = 3) and an *XPC*^KO^ ASC (n = 1) *in vitro.*

Similar to the *Ercc1^-/Δ^* mouse ASCs, loss of XPC in human ASCs induced an increase in the genome-wide number of base substitutions acquired per week. (Fig. 4C, Supplemental Table S5). In addition, the number of double base substitutions acquired per week was approximately 17 times higher (Fig. 4D, Supplemental Table S5, Supplemental Table S6). We did not observe a marked change in the genomic distribution of acquired mutations as a result of *XPC* deletion in human ASCs, nor a change in transcriptional strand bias (Supplemental Fig. S10C-D). In total, approximately 39% of the increase in base substitutions in the *XPC*^KO^ ASC can be explained by Signature 8 (Fig. 4E, Supplemental Fig. S10B), confirming that NER-deficiency can cause an increase in the number of Signature 8 mutations, independent of tissue-type or species.

### Detecting NER-deficiency in human breast cancers genomes

To identify whether NER-deficiency can be linked to an increase in Signature 8 mutations in human cancer as well, we looked into publicly available whole-genome sequencing data of 344 breast tumors (Nik-Zainal et al. 2016). Approximately 70% of these tumors have accumulated Signature 8 mutations (Nik-Zainal et al. 2016). NER-status was predicted by assessing the presence of protein-coding mutations and the copy number status of 66 NER-genes (Pearl et al. 2015). NER-deficient samples were defined as being hit by a biallelic loss-of-function mutation in at least one of the NER-genes. We excluded 274 samples with a mutation in one copy of any NER-gene from our analysis, as it is not possible to reliably predict the NER-status of these samples. By these definitions, we identified 27 NER-deficient samples and 43 NER-proficient samples.

NER-proficient and NER-deficient breast cancers have accumulated a median of 3,399 base substitutions (mean 3,968, standard deviation 2,708) and 4,368 base substitutions (mean 6,405, standard deviation 6,666) per sample, respectively (Supplemental Fig. S11A). To characterize whether NER-status affects the accumulation of Signature 8 mutations (http://cancer.sanger.ac.uk/cosmic/signatures), 96-channel mutational profiles of the somatic mutations were generated for all 344 breast tumors (Fig. 5A) and the contribution of each COSMIC mutational signature was assessed. 12 COSMIC mutational signatures contributed to <10% of the mutational profiles of all 344 tumors and were therefore excluded from subsequent analyses. The mutational profiles of the 70 breast tumors with predicted NER-status (Fig. 5A) were reconstructed using the remaining 18 mutational signatures. In line with previous observations, NER-deficient tumors have acquired 208 additional Signature 8 mutations in comparison to NER-proficient tumors (Fig. 5B; *P* = 0.02, Wilcoxon rank-sum test). Furthermore, Signature 8 has the largest effect size of all 18 COSMIC mutational signatures (Supplemental Fig. S11B).

**Figure 5.**
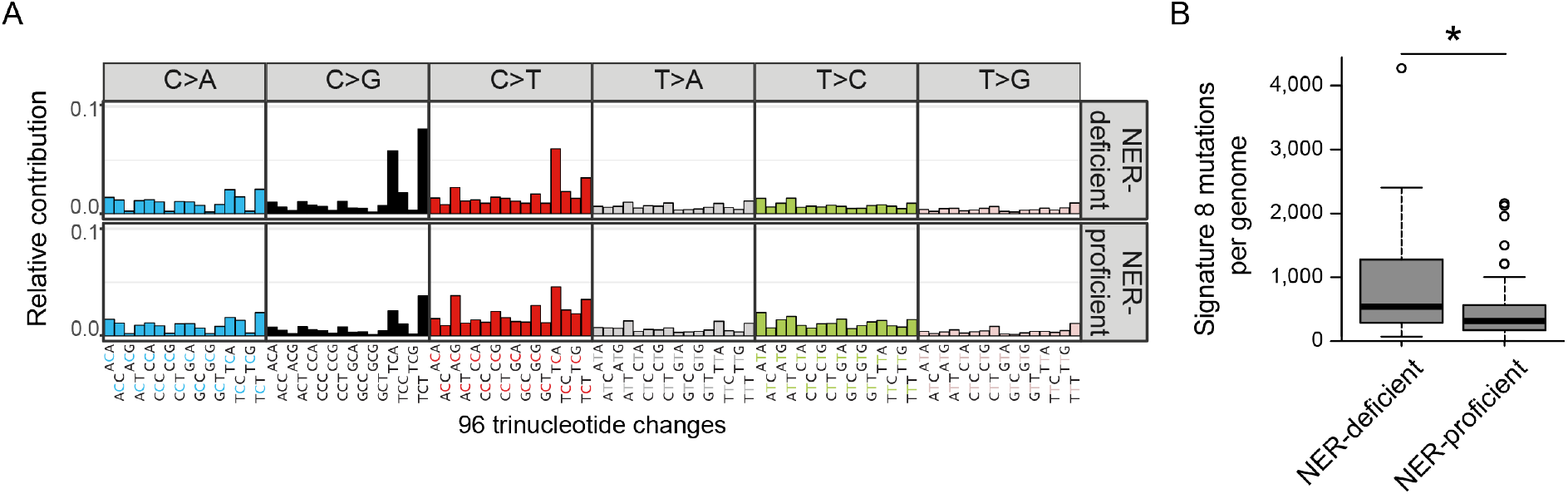
Mutation accumulation in predicted NER-deficient and NER-proficient breast cancer whole-genomes. (A) Relative contribution of each indicated context-dependent base substitution type to the average 96-channel mutational profiles of NER-deficient and NER-proficient breast cancer samples. (B) Number of Signature 8 mutations in NER-deficient and NER-proficient breast cancer whole-genomes (n = 27 and n = 43, respectively). Asterisk indicates significant difference (P < 0.05, Wilcoxon rank-sum test).

## DISCUSSION

We exploited mouse knockouts, organoid culturing, CRISPR-Cas9 gene-editing, WGS, and mutational signature analyses to study the genome-wide mutational consequences of NER-deficiency in individual ASCs of human and mice. Our results show that loss of ERCC1 induces a significant increase in the accumulation of base substitutions in liver ASCs, but not in small intestinal ASCs. Interestingly, the mutational increase coincides with the tissue-specific pathological aging phenotype observed in *Ercc1^-/Δ^* mice (Dollé et al. 2011; Gregg et al. 2012). A possible explanation for this difference between tissues is that liver ASCs might be more dependent on DNA repair facilitated by ERCC1 compared with small intestinal ASCs, e.g. as a result of tissue-specific mutagen exposure. In line with this, WT liver ASCs show a higher basal expression of *Ercc1* and other NER genes compared with WT small intestinal ASCs. However, the transcription levels of DNA repair components do not necessarily reflect DNA repair-activity, due to post-transcriptional regulation (Naipal et al. 2015). Alternatively, liver and small intestinal ASCs might cope differently with unrepaired DNA damage as a result of loss of ERCC1, such as the utilization of alternative DNA repair mechanisms, like translesion synthesis (TLS) polymerases, to bypass polymerase-blocking lesions, or differential induction of apoptosis or senescence.

ERCC1 is involved in multiple DNA repair pathways, including TC-NER, GG-NER, SSA, and crosslink repair. Previously, it has been shown that SSA- and crosslink repair-deficiencies result in increased number of indels and SVs in mice, whereas NER-deficiency introduces base substitutions (Dollé et al. 2006). Since we only observe an increase in base substitutions, NER-deficiency is likely responsible for the mutational consequences of loss of ERCC1 in liver ASCs *in vivo.* If TC-NER-deficiency underlies the differential mutation accumulation, this would be reflected by an increase in mutations in expressed genes in *Ercc1^-/Δ^* mice. However, WT and *Ercc1^-/Δ^* cells show a similar depletion of mutations in genes, indicating that the observed mutational consequences of impaired ERCC1 is rather an effect of defective GG-NER. In line with this, we show that GG-NER-deficiency can also induce an increase in the number of base substitutions in a human small intestinal organoid culture that is deleted for GG-NER component *XPC.* More specifically, the increased base substitution load can be largely explained by an increased contribution of Signature 8 in both systems. In line with our observations, a mutational signature similar to Signature 8 has been shown to increase with age in the neurons of NER-deficient patients (Lodato et al. 2017).

Until now, the etiology of Signature 8 was unknown (http://cancer.sanger.ac.uk/cosmic/signatures). As Signature 8 mutations are also detected in healthy human and mouse ASCs (Fig. 3C, Fig. 4E), this signature most likely represents a mutagenic process that is generally active in normal cells and not repaired 100% effective. Signature 8 is characterized by C:G > A:T mutations and is associated with double base substitutions (Alexandrov et al. 2013; Nik-Zainal et al. 2016). C:G > A:T mutations have been linked to several processes, including oxidative stress (Kamiya et al. 1995; Degtyareva et al. 2013). Consistently, organoid culturing causes mutations indicative of high oxidative stress (Blokzijl et al. 2016). Interestingly, NER has been suggested to play a role in the repair of tandem DNA lesions that result from oxidative stress (Bergeron et al. 2010; Cadet et al. 2012). If left unrepaired, these lesions can block regular DNA polymerases, but can be bypassed by error-prone TLS polymerases, resulting in increased incorporation of tandem mutations (Cadet et al. 2012). Moreover, it has been shown that oxidative stress results in increased induction of double base substitutions in NER-deficient human fibroblasts (Lee 2002). In line with this, we observe a significant increase in the double base substitution load in mouse liver ASCs and a similar trend in the human ASC culture as a result of NER-deficiency, yet the number of double base substitutions is much lower than single base substitutions. Thus Signature 8 could reflect oxidative DNA damage bypassed by TLS.

Although NER-deficiency does not affect the base substitution load in the mouse small intestine, it does result in an increased contribution of Signature 8 mutations. This is in clear contrast to mouse liver ASCs, where NER-deficiency has both a qualitative and quantitative consequence on the accumulation of base substitutions. More specifically, the absolute contribution of Signature 8 mutations is similar in WT liver and *Ercc1^-/Δ^* small intestinal ASCs. This clearly demonstrates that DNA-repair deficiency can have tissue-specific consequences, but also indicates that the absolute contribution of Signature 8 mutations should be compared to the basal contribution in the same tissue in order to detect NER-deficiency.

We did not observe a notable contribution of signatures that have been previously observed in liver cancer in ASCs of *Ercc1^-/Δ^* livers (http://cancer.sanger.ac.uk/cosmic/signatures) (Supplemental Fig. S6B). This finding suggests that the mutational processes that underlie these signatures are only active after oncogenic transformation, or that mutagen exposure in liver cancer (progenitor) cells is different from *in vivo* mouse ASCs and *in vitro* human ASCs. Liver cancer-specific Signature 24, for example, is associated with Aflatoxin intake (Huang et al. 2017), a substance to which our mice and organoids were not exposed. In addition, Signature 1 and Signature 5, which have been previously associated with age (Blokzijl et al. 2016; Alexandrov et al. 2015), did not have an increased contribution in the ASCs of progeroid *Ercc1^-/Δ^* mice. Finally, a high contribution of mutational Signature 5 has been linked to the presence of somatic mutations in *ERCC2,* a key factor in both TC-NER and GG-NER, in human urothelial cancer (Kim et al. 2016; Iyama and Wilson 2013). As mentioned however, we did not observe an increase in Signature 5 contribution in the ASCs without ERCC1 or XPC. This discrepancy in mutational consequences could reflect various differences between these systems, such as different effects of the mutations on protein function, distinct roles of the proteins, or tumor-and/or tissue-specific activity of mutagenic damage and/or DNA repair processes. In our study, we deleted specific NER components in an otherwise normal genetic background, providing us with the unique opportunity to directly characterize the mutational consequences of NER-deficiency.

The challenge of coupling mutational signatures to mutational processes based on genome sequencing data of tumors is illustrated by our analyses of the breast cancer genomes. As the number of mutations attributed to a signature typically increases at a higher mutational load and the mutational loads differ greatly between tumor types (Alexandrov et al. 2013), it is important to compare signature contributions between samples within a single tumor type. However, the majority of the breast cancer samples (~80%) carried a single mutation in a NER-gene and since the other copy might be inactivated through e.g. epigenetic silencing, these samples were excluded from the analysis. This resulted in a low sample size. Nonetheless, genomes of NER-deficient breast cancer patients show a higher number of Signature 8 mutations, which is in line with our observations in the ASCs. Further optimization of mutational signature definitions may aid to discriminate NER-deficient from NER-proficient tumors fully.

Determination of the NER-capacity of tumors can be important for precision medicine, as it has been shown that tumors with mutations in NER genes (Stubbert et al. 2010; Van Allen et al. 2014; Amable 2016; Zhang et al. 2017), and tumors with low expression of *ERCC1* (Olaussen et al. 2006; Li et al. 2000; Amable 2016) are sensitive to cisplatin treatment. However, translation of these findings into the clinical setting has been challenging, because connecting tumor biopsy mRNA levels and immunohistochemistry measurements to NER-activity remains an unresolved issue (Bowden 2014), and interpreting the effects of mutations in DNA repair genes on NER-capacity is challenging. Rather than looking for the presence of causal events, mutational catalogs can be used as a functional readout of NER-capacity in tumors. Here, we show that NER-deficiency can induce an increase in Signature 8 mutations in both mouse and human ASCs. Signature 8 is found in medulloblastoma, bladder cancer, bone cancer, lung squamous cell carcinoma, ovarian cancer, pancreatic cancer and prostate cancer (Alexandrov et al. 2013, 2018). Furthermore, Signature 8 contributes to the mutational profile of the majority of breast cancer tumors (Nik-Zainal et al. 2016; Alexandrov et al. 2013). Our results show that, besides the mutational status of NER genes, an increase in the number of Signature 8 mutations with respect to the normal number of Signature 8 mutations in a cancer type might serve as a novel biomarker for (GG-)NER-deficiency and has the potential to guide treatment decision. Clinical studies will be required to demonstrate the added predictive value of Signature 8 mutations for NER-deficiency and/or treatment response.

## METHODS

### Mouse tissue material

*Ercc1^-/Δ^* mice were generated and maintained as previously described (Vermeij et al., 2016). Briefly, by crossing *Ercc1^Δ/+^* (C57BL6J or FVB background) with *Ercc1^+/−^* mice (FVB or C57BL6J background), *Ercc1^-/Δ^* mice were generated in a uniform F1 C57BL6J/FVB hybrid background. Wild type F1 littermates were used as controls. Animals were housed in individually ventilated cages under specific pathogen-free conditions in a controlled environment (20–22 °C, 12 h light : 12 h dark cycle). Experiments were performed in accordance with the Principles of Laboratory Animal Care and with the guidelines approved by the Dutch Ethical Committee in full accordance with European legislation.

We used three 15-week old female *Ercc1^-/Δ^* mice and three female WT littermates for our experiments. Tails were harvested and stored at -20°C. Livers and small intestines were harvested and kept on ice in Adv+++ medium (Advanced DMEM/F-12 with 1% GlutaMAX, HEPES 10 mM and 1% penicillin/streptomycin) for a few hours until further processing.

### Human tissue material

Endoscopic biopsies were performed at the University Medical Center Utrecht and the Wilhelmina Children’s Hospital. The patients’ informed consent was obtained and this study was approved by the ethical committee of University Medical Center Utrecht.

### Generation of clonal *Ercc1^-/Δ^* and WT mouse organoid cultures

Single liver ASCs were isolated from livers as described previously (Kuijk et al. 2016). Liver organoid cultures were initiated by culturing the liver ASCs in BME overlaid with mouse liver culture initiation medium (50% Adv+++ medium, 35% WNT3A conditioned medium (produced in house), 5% NOGGIN conditioned medium (produced in house), 5% RSPOI conditioned medium (produced in house), 1x B27 without retinoic acid, 1x N2, 1x Primocin, 10mM Nicotinamide, 0.625mM N-acetylcysteine, 100ng/ml FGF-10, 10μM ROCKi, 50 ng/ml HGF, 10nM Gastrin, and 50ng/ml hEGF). 1.5 week after culture initiation, clonal organoid liver cultures were generated and expanded according to protocol (Jager et al. 2018) in mouse liver expansion medium (90% Adv+++ medium, 5% RSPOI conditioned medium (produced in house), 1x B27 without retinoic acid, 1x N2, 1x Primocin, 10mM Nicotinamide, 0. 625mM N-acetylcysteine, 100ng/ml FGF-10, 50 ng/ml HGF, 10nM Gastrin, and 50ng/ml hEGF).

Crypts were isolated from small intestines as described previously (Sato et al. 2009). Small intestinal organoid cultures were initiated by culturing the small intestinal ASCs in matrigel overlaid with mouse small intestine medium (50% WNT3A conditioned medium (produced in house), 30% Adv+++ medium, 10% NOGGIN conditioned medium (produced in house), 10% RSPOI conditioned medium (produced in house), 1x B27, 1x hES Cell Cloning & Recovery Supplement, 1x Primocin, 10μM ROCKi, 1.25mM N-acetylcysteine, and 50ng/ml hEGF). Clonal small intestinal organoid cultures were generated by picking single organoids manually and clonally expanding these organoid cultures according to protocol in mouse small intestine medium (Jager et al. 2018). Culture expansion failed for the small intestine of mouse WT1.

### Generation of a clonal and subclonal *XPC*^KO^ organoid culture

Clonal *XPC*^KO^ organoid cultures were generated from a small intestinal bulk organoid culture derived previously (Blokzijl et al. 2016) using the CRISPR-Cas9 gene-editing technique as described in (Drost et al. 2017). One clonal human *XPC*^KO^ organoid culture was obtained and cultured for 72 days in human small intestinal organoid medium (50% WNT3A conditioned medium (produced in house), 30% Adv+++ medium, 20% RSPOI conditioned medium (produced in house), 1x B27, 1x Primocin, 1.25mM N-acetylcysteine, 0.5μM A83-1, 10μM SB202190, 100ng/ml recombinant Noggin, and 50ng/ml hEGF). Subsequently, a subclonal culture was derived according to protocol (Jager et al. 2018).

### Western blot

Protein samples from mouse organoid cultures were collected in Laemmli buffer and measured using the Qubit^tm^ 3.0 Fluorometer (Thermo Fisher Scientific) with the Qubit^tm^ Protein Assay Kit (Thermo Fisher Scientific, Q33211). Protein samples from human organoid cultures were collected in Laemmli buffer and measured using a Lowry protein assay. 30μg of protein per sample was run on a 10% SDS page gel. Subsequently, the proteins were transferred to a nitrocellulose membrane. After transfer, the membrane was blocked for 1 hour using 5% ELK (Campina) at room temperature and subsequently incubated overnight with the primary antibody (ERCC1: Abcam, ab129267; XPC: Cell Signaling Technology; #12701). Secondary antibody was incubated 1 hour at room temperature, and subsequently proteins were visualized using the Amersham ECL Western blotting analysis system (GE Healthcare, RPN2109) and the Amersham Imager 600 system (GE Healthcare).

### RNA sequencing and differential expression analysis of *Ercc1^-/Δ^* and WT mouse organoid cultures

For each mouse (three *Ercc1^-/Δ^* mice and three WT littermates), we performed RNA sequencing on one clonal organoid culture from the liver and the small intestine. An additional small intestinal organoid clone was sequenced of mice WT2 and WT3 to increase the amount of replicates for differential expression analysis, as culture expansion failed for the small intestine of WT1. Total RNA was collected in TRIzol and purified from all organoid cultures using the Qiasymphony (Qiagen). RNA libraries for Illumina sequencing were generated from 50 ng of poly-A selected mRNA using the Neoprep (Illumina) and sequenced 2 x 75 bp paired-end to approximately 3300 Million base pairs per sample with the Illumina NextSeq 500 at the Utrecht Sequencing Facility.

RNA sequencing reads were mapped with STAR v.2.4.2a to the mouse reference genome GRCm38. The BAM files were sorted with Sambamba v0.5.8 and reads were counted with HTSeq-count version 0.6.1p1 (default settings) to exons as defined in GRCm38v70.gtf (Ensembl). Non-uniquely mapped reads were not counted. Subsequently, DESeq v1.28.0 was used to normalize counts. DESeq nbinomTest was used to test for differential expression (1) of *Ercc1* between *Ercc1^-/Δ^* and WT liver ASCs, (2) of *Ercc1* between *Ercc1^-/Δ^* and WT small intestinal ASCs, (3) of 83 other DNA repair genes (Casorelli et al. 2006) between *Ercc1^-/Δ^* and WT liver ASCs, and (4) between *Ercc1^-/Δ^* and WT small intestinal ASCs, and (5) of 9 NER genes between the WT liver and WT small intestinal ASCs. Differentially expressed genes with *q* < 0.05 (Benjamini-Hochberg FDR multiple-testing correction) were considered significant.

### WGS and read alignment

DNA was isolated from mouse liver organoid cultures and mouse control (tail) samples using the genomic tip 20-G kit (Qiagen) and from mouse small intestinal organoid samples and the human *XPC*^KO^ sample using the Qiasymphony (Qiagen). DNA libraries for Illumina sequencing were generated from 200 ng genomic DNA using standard protocols (Illumina) and sequenced 2 x 100 bp paired-end to 30X base coverage with the Illumina HiSeq Xten at the Hartwig Medical Foundation. The sequence reads of *XPC*^KO^ were mapped to the GRCh37 human reference genome using using the Burrows–Wheeler Aligner (BWA) v0.7.5a (Li and Durbin 2009), with settings ‘-t 4 -c 100 -M’. The mapped data of clonal *XPCA*^WT^ organoids was previously generated in the study (‘donor_id’ 6) (Blokzijl et al. 2016). The sequence reads of the mouse ASCs were mapped to the GRCm38 mouse reference genome using using the Burrows–Wheeler Aligner (BWA) v0.7.5a (Li and Durbin 2009), with settings ‘-t 4 -c 100 -M’. The WGS data of the tails confirmed that the *Ercc1^-/Δ^* mice have compound heterozygous mutations in *Ercc1* and the WT littermates do not (Supplemental Fig. S12).

### Callable genome

The callable genome was defined for all sequenced samples using the GATK CallableLoci tool v3.4.46 (Van der Auwera et al. 2013) with default settings and additional optional parameters ‘minBaseQuality 10’, ‘minMappingQuality 10’, ‘maxFractionOfReadsWithLowMAPQ 20’, and ‘minDepth 20’. ‘CALLABLE’ regions were extracted from every output file. Subsequently, genomic regions that were callable (1) in the mouse organoid clone and the control (tail) sample, and (2) in the human organoid clone, subclone, and control (blood) were intersected to define a genomic region that is surveyed in all samples that were compared. Approximately 90 ± 1% of the autosomal genome was surveyed in every mouse clone (Supplemental Table S2), and 73 - 88% of the autosomal genome was surveyed in each human subclone (Supplemental Table S5).

### Base substitution and indel calling

For both human and mouse samples, base substitutions and indels were multi-sample called with GATK HaplotypeCaller v3.4.46 with default settings and additional options ‘-stand_call_conf 30 -stand_emit_conf 15’ and GATK Queue v3.4.46. For mouse samples the quality of the calls was assessed using GATK VariantFiltration v3.4.46 with options ‘QD < 2.0, MQ < 40.0, FS > 60.0, HaplotypeScore > 13.0, MQRankSum < -12.5, ReadPosRankSum < -8.0’ for base substitutions and ‘QD < 2.0, FS > 200.0, ReadPosRankSum < -20.0’ for indels, with additional options ‘clusterSize 3’ and ‘clusterWindowSize 35’. For human samples the quality of the calls was assessed using GATK VariantFiltration v3.4.46 with options ‘QD < 2.0, MQ < 40.0, FS > 60.0, HaplotypeScore > 13.0, MQRankSum < -12.5, ReadPosRankSum < -8.0, MQ0 >= 4 && ((MQ0 / (1.0 * DP)) > 0.1), DP < 5, QUAL < 30, QUAL >= 30.0 && QUAL < 50.0, SOR > 4.0’ for base substitutions and ‘QD < 2.0, FS > 200.0, ReadPosRankSum < -20.0, MQ0 >= 4 && ((MQ0 / (1.0 * DP)) > 0.1), DP < 5, QUAL < 30.0, QUAL >= 30.0 && QUAL < 50.0, SOR > 10.0’ for indels, with additional options ‘clusterSize 3’ and ‘clusterWindowSize 10.

### Base substitution filtering

To obtain high-quality catalogs of somatic base substitutions, we applied a comprehensive filtering procedure. For the mouse samples, we only considered positions on the autosomal genome that were callable (see “Callable genome”) in both the organoid and control (tail) sample. We excluded positions at which indels were called, as these positions likely represent false-positive base substitution calls. Furthermore, we only included positions with a ‘PASS’ flag by GATK VariantFiltration, a GATK phred-scaled quality score ≥ 100, a sample-level genotype quality of 99 in the organoid culture and ≥ 10 in the control (tail) sample, and a coverage of ≥ 20X in the organoid and the tail sample. We subsequently excluded variants with any evidence in another organoid sample or control (tail) sample of the same mouse to remove germline variants. To exclude potentially missed germline events, we also removed positions that have any evidence in the organoid and/or control samples of the other mice. Finally, we excluded positions with a variant allele frequency (VAF) < 0.3 in the organoid sample to exclude mutations that were induced after the clonal step.

For the human samples, we only considered positions on the autosomal genome that were callable (see “Callable genome”) in the control (blood) sample, clonal organoid and subclonal organoid culture. We considered mutations with a ‘PASS’ flag by GATK VariantFiltration and a GATK phred-scaled quality score ≥ 100. For both the clonal and subclonal organoid cultures, all variants with evidence in the control (blood) sample were excluded, to remove germline variants. To exclude potentially missed germline events, we removed positions that are in the Single Nucleotide Polymorphism Database v137.b3730, or in a blacklist with positions that are recurrent in unmatched individuals (BED-file available upon request). Subsequently, for both the clonal and subclonal cultures, all variants with a VAF < 0.3 were excluded. Finally, the resulting somatic base substitution catalogs of the clonal and subclonal cultures were compared and all events unique to the subclonal organoid were considered to be accumulated after the XPC deletion, that is: between the two sequential clonal expansion steps.

### Validation of base substitutions in *Ercc1^-/Δ^* and WT mouse organoid cultures

To independently validate all base substitution positions, new sequencing libraries were generated from DNA samples of all 11 mouse organoid cultures and of the tail of WT1 (DNA samples were isolated in ‘WGS and read alignment’) using the Twist Human Core Exome v1.3 Complete Kit. The libraries were pooled and size-selected twice using 0.55X and 0.8X AMPure XP beads. For all base substitutions, an enrichment probe of 120 bp was designed for both the reference and variant allele with a minimum number of repeats and with the base substitution position at least 10 bp from the end. The probes were produced by Twist Bioscience. Subsequently, the pooled libraries were enriched with the enrichment probes in two enrichment reactions using the Twist Human Core Exome v1.3 Complete Kit and sequenced 2 x 150 bp paired-end with the Illumina NextSeq 500 at the Utrecht Sequencing Facility. Base substitutions were called as described in ‘Base substitution and indel calling’. Variants were considered true if they were called with a filter ‘PASS’ in 1 out of 12 samples. The remaining 220 variants were checked manually in IGV and considered true if they were found in 1 out of 12 samples at an allele frequency of > 10%. In total, 4,130/4,238 variants (97.5%) were confirmed using this approach (Supplemental file S2).

### Clonality of organoid cultures

We validated whether the organoid samples were clonal based on the VAF of somatic base substitutions, before the final filter step (VAF < 0.3). Each cell acquires its own set of somatic mutations and the reads supporting a mutation will be diluted in the WGS data of non-clonal samples, resulting in a low VAF. After extensive filtering of somatic base substitutions, liver organoid samples from WT1, WT2, and *Ercc1^-/Δ^*2 showed a shift in the VAF-peak away from 0.5 and therefore these samples were excluded from further analyses (Supplemental Fig. S13). An additional liver organoid culture from these mice was sequenced and these samples were confirmed to be clonal (Supplemental Fig. S13).

### Double base substitutions

We selected base substitutions from the filtered variant call format (VCF) files that were called on consecutive bases in the mouse or human reference genome. The double base substitutions were subsequently manually checked in the Integrative Genomics Viewer (IGV) to exclude double base substitutions present in the control sample, and/or with many base substitutions or indels in the region, as these are (likely) false positives.

### Indel filtration of *Ercc1^-/Δ^* and WT mouse organoid cultures

We only considered positions on the autosomal genome that were callable (see “Callable genome”) and had a sequencing depth of ≥ 20X in both the organoid sample and the control (tail) sample. We excluded positions that overlap with a base substitution. Furthermore, we only considered positions with a filter ‘PASS’ from VariantFiltration, a GATK phred-scaled quality score > 250 and a sample-level genotype quality of 99 in both the organoid sample and the control (tail) sample. We subsequently excluded Indels that are located within 50 base pairs of an indel called in another organoid sample and indels with any evidence in another organoid sample or a control (tail) sample. Finally, we excluded positions with a VAF < 0.3 in the organoid sample.

### SV calling and filtration of *Ercc1^-/Δ^* and WT mouse organoid cultures

SVs were called with DELLY v0.7.2 with settings ‘type DEL DUP INV TRA INS’, ‘map-qual 1’, ‘mad-cutoff 9’, ‘min-flank 13’, and ‘geno-qual 5’ (Rausch et al. 2012). We only considered SVs of at least 100 bp on the autosomal chromosomes that were called with a filter ‘PASS’, and a sample-specific genotype quality of at least 90 in the organoid culture and the control sample. We subsequently excluded positions with any evidence in the control (tail) sample. The filtered SVs were finally checked manually in IGV to reduce false-positives and we excluded SVs present in the tail sample, with no visible change in the read-depth (for duplications and deletions), and/or with many base substitutions in the region.

### Genome-wide copy number profiles of *Ercc1^-/Δ^* and WT mouse organoid cultures

To generate a virtual karyotype, genome-wide copy number states were determined using FreeC v7.2 with settings ‘ploidy 2’, ‘window 1000’ and ‘telocentromeric 50000’ (Boeva et al. 2012). Subsequently, the average copy number across bins of 500,000 bp was calculated and plotted to assess genome stability.

### Base substitution types

We retrieved the base substitution types from all the filtered VCF files, converted them to the 6 types of base substitutions that are distinguished by convention, and generated a mutation spectrum (the C>T changes at NpCpG sites are considered separately from C>T changes at other sites) for the four ASC groups (*Ercc1^-/Δ^* liver, *Ercc1^-/Δ^* small intestine, WT liver, and WT small intestine), as well as *XPC*^KO^, *XPCA*^WT^1, *XPCA*^WT^2, and *XPCA*^WT^3 ASCs. X^2^-tests were performed to determine whether the mutation spectra differ significantly between (1) mouse WT and *Ercc1^-/Δ^* liver ASCs, and (2) mouse WT and *Ercc1^-/Δ^* small intestinal ASCs. *P* values were corrected for multiple testing using Benjamini-Hochberg FDR correction, and differences in mutation rates between *Ercc1^-/Δ^* and WT mouse ASCs with *q* < 0.05 were considered significant.

We retrieved the sequence context for all base substitutions to generate the 96- channel mutational profiles for each assessed ASC. Subsequently, the centroid of the 96- channel mutational profiles was calculated per mouse ASC group. Pairwise cosine similarities of all 96-channel mutational profiles and of all centroids were computed. We also calculated the cosine similarities of the 96-channel mutational profiles and centroids with all 30 COSMIC mutational signatures (http://cancer.sanger.ac.uk/cosmic/signatures) (Supplemental Fig. S7). These analyses were performed with the R package MutationalPatterns (Blokzijl et al. 2018).

### *De novo* mutational signature extraction

We extracted two signatures using non-negative matrix factorization (NMF) from the 96- channel mutational profiles of the mouse ASCs. Although the number of base substitutions is low for this dimension reduction approach, it does provide an unbiased method to characterize the mutational processes that have been active in the ASCs. Subsequently, we computed the absolute contribution of these *de novo* extracted signatures to the centroids of the mouse ASC groups. We also calculated the cosine similarity of these two mutational signatures to the 30 COSMIC mutational signatures (http://cancer.sanger.ac.uk/cosmic/signatures) and to the 96-channel centroid of six small intestinal ASCs from two old mice that was published previously (Behjati et al. 2014). These analyses were performed with MutationalPatterns (Blokzijl et al. 2018).

### Quantification of the contribution of C?SMIC mutational signatures to the 96-channel mutational profiles

We estimated the contribution of the 30 COSMIC mutational signatures (http://cancer.sanger.ac.uk/cosmic/signatures) to the centroids of each mouse ASC group and to the 96-channel mutational profiles of the human organoids using MutationalPatterns (Blokzijl et al. 2018) (Supplemental Fig. S6B, Supplemental Fig. S10B). We ranked the COSMIC signatures based on the total contribution of these signatures to the centroids of the mouse samples. Next, we iteratively reconstructed the centroids of the ASC groups, first using the top 2 COSMIC signatures, and in each iteration the next COSMIC signature was included until all 30 signatures were used. The cosine similarity was calculated between the original and the reconstructed centroid for each mouse ASC group (Supplemental Fig. S6A). As expected, the addition of more signatures increases the similarity of the reconstructed centroids with the original centroids, but after 10 COSMIC signatures the cosine similarities plateau (Supplemental Fig. S6A). Therefore, we used the signature contribution with this subset of 10 COSMIC signatures to the centroids of the four ASC groups (Fig. 3B-C).

### Determination of the statistical significance of differences in signature contributions

A bootstrap resampling - similar to that performed in (Zou et al. 2018) - was applied to generate 7,000 replicas of the 96-channel mutational profile of each WT liver ASC (n = 3), which yielded 21,000 WT liver replicas in total. Subsequently, 3 replicas were randomly selected and the relative contribution of 30 COSMIC signatures was determined for their centroid. Euclidean distance *d_WT_* was calculated between the relative signature contributions of the replicas centroid and that of the original centroid. This was repeated 10,000 times to construct a distribution of *d_WT_* (Supplemental Figure 6C). Next, the threshold distance with *P* value = 0.05, *d_WT_0.05_,* was identified. The same approach was taken to generate 7,000 replicas of each *Ercc1^-/Δ^* (MUT) liver ASC (n = 3) and construct a distribution of *d_MUT_* (Supplemental Figure 6C). The Euclidean distance *d* between the relative signature contributions of the original WT and *Ercc1^-/Δ^* liver centroids were considered to be significantly different when *d* > *d_MUT_* and d > d_WT_. Similarly, bootstrap distributions were generated for WT and *Ercc1^-/Δ^* (MUT) small intestine (Supplemental Figure 6D), with the exception that for the generation of the *d_MUT_* distribution only 2 replicas were randomly selected in each permutation, as there are only 2 WT small intestinal ASC samples in the original set. Finally, we repeated the same analyses for the relative contributions of the subset of 10 COSMIC signatures for both liver (Supplemental Figure 6E) and small intestine (Supplemental Figure 6F).

### Enrichment or depletion of base substitutions in genomic regions

To test whether the base substitutions appear more or less frequently than expected in genes, promoters, promoter-flanking, and enhancer regions, we loaded the UCSC Known Genes tables as TxDb objects for Mm10 (Team BC and Maintainer 2016) and Hg19 (Carlson and Maintainer 2015), and the regulatory features for Mm10 and Hg19 from Ensembl using biomaRt (Durinck et al. 2005, 2009). We tested for enrichment or depletion of base substitutions in the genomic regions per ASC group (*Ercc1^-/Δ^* liver, *Ercc1^-/Δ^* small intestine, WT liver, WT small intestine, *XPC*^KO^ and *XPCA*^WT^) using a one-sided Binomial test with MutationalPatterns (Blokzijl et al. 2018), which corrects for the surveyed genomic areas (Supplemental Fig. S9A, Supplemental Fig. S10C). Two-sided Poisson tests were performed to test for significant differences in the ratio of base substitutions within a genomic region divided by the total number of base substitutions between (1) mouse WT and *Ercc1^-/Δ^* liver ASCs and (2) mouse WT and *Ercc1^-/Δ^* small intestinal ASCs (Supplemental Fig. S9A). Differences in mutation rates with *q* < 0.05 (Benjamini-Hochberg FDR multiple-testing correction) were considered significant.

To test whether base substitutions occur more frequently in more highly expressed genes in the NER-deficient mouse ASCs, we first selected base substitutions that occurred within genes in the mouse ASCs. Per ASC group, we next determined the average Reads Per Kilobase per Million mapped reads (RPKM) of these genes. Two-sided ř-tests were performed to test for significant difference in the average expression of genes that carry a somatic mutation between (1) mouse WT and *Ercc1^-/Δ^* liver ASCs, and (2) mouse WT and *Ercc1^-/Δ^* small intestinal ASCs (Supplemental Fig. S9B). Differences in gene expression distributions with *q* < 0.05 (Benjamini-Hochberg FDR multiple-testing correction) were considered significant.

### Transcriptional strand bias of base substitutions

For the base substitutions within genes we determined whether the mutations are located on the transcribed or the non-transcribed strand. To this end, we determined whether the mutated “C” or “T” base is on the same strand as the gene definition, which is untranscribed, or the opposite strand, which is transcribed. We generated a 192-channel mutational profile per ASC group with the relative contribution of each mutation type with separate bars for the mutations on the transcribed and untranscribed strand, and calculated the significance of the strand bias using a two-sided Poisson test with MutationalPatterns (Supplemental Fig. S9C, Supplemental Fig. S10D) (Blokzijl et al. 2018). Furthermore, we performed two-sided Poisson tests to test whether there is a significant difference in strand bias per mutation type between (1) mouse WT and *Ercc1^-/Δ^* liver ASCs and (2) mouse WT and *Ercc1^-/Δ^* small intestinal ASCs (Supplemental Fig. S9C). Differences in strand bias with an adjusted P-value *q* < 0.05 (Benjamini-Hochberg FDR multiple-testing correction) were considered significant.

### Calculation and comparison of mutation rates

To calculate the mutation rates per genome per week, we quantified the number of somatic base substitutions, double nucleotide mutations, indels, and SVs for each mouse ASC. Moreover, we quantified the number of base substitutions, double base substitutions and Signature 8 mutations for the human ASCs. All event counts were extrapolated to the entire autosomal genome using the callable genome length (see” Callable genome”) for both mouse and human ASCs to correct for differences in the surveyed genome. Subsequently, the mutation rates were calculated by dividing the extrapolated number of mutations by the number of weeks in which the mutations were accumulated (WT and *Ercc1^-/Δ^* mouse organoids: 16 weeks (15 weeks during life and 1 week *in vitro*); *XPCA*^WT^ human organoids: 20.6 weeks; *XPC*^KO^ human organoids 10.3 weeks). To determine the proportion of additionally accumulated mutations in the *XPC*^KO^ culture that can be attributed to Signature 8 in human ASCs, we first calculated the increase in base substitutions and the increase in Signature 8 mutations of *XPC*^KO^ compared to *XPCA*^WT^1, *XPCA*^WT^2, and *XPCA*^WT^3 separately. We then divided the increase in Signature 8 mutations by the total increase in base substitutions.

Two-tailed *t*-tests were performed to determine whether the mutation rates differ significantly between (1) mouse WT and *Ercc1^-/Δ^* liver ASCs, and (2) mouse WT and *Ercc1^-/Δ^* small intestinal ASCs. Of note, these tests assume that the data is normally distributed. Differences in mutation rates between *Ercc1^-/Δ^* and WT mouse ASCs with *q* < 0.05 (Benjamini-Hochberg FDR multiple-testing correction) were considered significant.

### Analysis of mutational patterns and signatures in breast cancer whole-genome sequences

344 breast cancer samples with publicly available SNV, indels, and CNV calls obtained from tumor-normal samples were included in the analysis (Nik-Zainal et al, 2016). Samples with a biallelic inactivation (biallelic deletion, biallelic nonsense, splice site, nonsynonymous mutation or frameshift indel, or two or more independent mutations of these types) of at least one NER-related gene (66 genes, (Pearl et al. 2015); GTF2H5 was excluded because of missing CNV calls) are considered as NER-deficient. Samples with no copy number depletions and no variants other than intronic SNVs and indels in any of the 66 NER-related genes are considered as NER-proficient. The remaining 274 samples are considered as having unknown NER-ability.

The number of base substitutions was extracted from each VCF file and a Wilcoxon rank-sum test was performed to determine whether the number of base substitutions is different between NER-proficient and NER-deficient samples. The 96-channel mutational profile of each sample was generated as described in the section “Base substitution types”. Subsequently, the 96-channel mutational profile of each sample was reconstructed using the 30 mutational signatures from COSMIC, as described in the section “Quantification of the contribution of COSMIC mutational signatures to the 96-channel mutational profiles”. Signatures with a contribution of < 10% in all 344 samples were excluded (signatures 4, 7, 10, 11, 14, 15, 22-25, 27, 28), and the 96-channel mutational profiles were finally reconstructed using the remaining 18 signatures. The cosine similarity between the observed 96-channel mutational profile and the reconstructed profile was above 0.95 for all samples, which indicates a very good fit of the signatures.

Based on this, the number of mutations per signature was estimated for each sample. Then, for each signature, the number of mutations was compared between the NER-deficient and NER-proficient samples using the median difference (the median of all pairwise differences between NER-deficient and NER-proficient samples). A Wilcoxon rank-sum test was performed to determine whether the number of Signature 8 mutations differs significantly between NER-deficient and NER-proficient breast tumors. Signature 20 is excluded from the analysis, because none of the NER-deficient or NER-proficient samples have a contribution of Signature 20.

## DATA ACCESS

The sequencing data of the mouse samples have been deposited at the European Nucleotide Archive under accession number ERP021379. The sequencing data of the human samples have been deposited at the European Genome-Phenome archive under accession numbers EGAS00001001682 and EGAS00001002681. Filtered VCF files are freely available at https://wgs11.op.umcutrecht.nl/NERdeficiency/

## CODE AVAILABILITY

All analysis scripts are available at https://github.com/UMCUGenetics/NER-deficiency.git or https://github.com/johannabertl/BRCA_DNA_repair.

## ACKNOWLEDGEMENTS

The authors would like to thank the the animal caretakers of the Erasmus MC for taking care of the mice and the Utrecht Sequencing Facility for providing the sequencing service and data. Utrecht Sequencing Facility is subsidized by the University Medical Center Utrecht, Hubrecht Institute and Utrecht University. This study was financially supported by the NWO Zwaartekracht program Cancer Genomics.nl.

## AUTHOR CONTRIBUTIONS

M.J., E.K., M.V., N.B., and R.B. performed organoid culturing. N.B. and R.B. generated western blots and sequenced the organoid cultures. M.J., F.B., J.B., R.J., S.B., J.L., and R.B. performed bioinformatic analyses. M.J., F.B., E.K., J.S.P., J.H., J.P., R.B., and E.C. were involved in the conceptual design of this study. M.J., F.B., R.B., and E.C. wrote the manuscript.

